# Keystone mutualists can facilitate transition between alternative ecosystem states in the soil

**DOI:** 10.1101/392993

**Authors:** Marie Duhamel, Joe Wan, Laura M. Bogar, R. Max Segnitz, Nora C. Duncritts, Kabir G. Peay

## Abstract

Symbioses between plants and microbial organisms can fundamentally alter the structure of ecosystems, from their species diversity to rates of nutrient cycling. Yet, many aspects of how differences in the prevalence of microbial symbioses arise are unclear. This is a key knowledge gap, as if co-variation in plant and microbial distributions are primarily determined by extrinsic abiotic factors then symbioses should exert little independent control over ecosystems. To examine the potential for alternative symbiotic communities to arise under similar conditions we examined biogeochemical cycling and microbial community structure in a coastal landscape where historical patterns of vegetation transition are known, allowing us to eliminate abiotic determinism. We found that alternative states in microbial community structure and ecosystem processes emerged under different plant species. Greenhouse studies further demonstrated that plant selection of symbiotic microbes is central to emergence of these alternative states and occurs independent of soil abiotic conditions. Moreover, we provide evidence that transition between states may be highly dependent on the presence of a small set of ruderal symbionts that are rare in mature systems but may act as keystone mutualists. Because differences between these alternative states can be directly linked to plant-microbe symbioses, independent of initial conditions, our results suggesting that biotic feedbacks between keystone symbiotic microbes and plants play a foundational role in the diversity and function of soils.

## Introduction

Early ecologists viewed community assembly as an organized process where species facilitate each other in progression towards a stable, climax state (Clements 1936). This “organismic” view of communities contrasts starkly with the individualistic theory of ecology (Gleason 1926), where communities are simply the local product of individual species’ responses to environmental conditions and probabilistic dispersal events. While Clements’ viewpoint has been largely abandoned by ecologists (McIntosh 1998), recent work on the microbiome suggests that symbioses and positive interactions are more common than previously expected (Gilbert et al. 2012, Peay 2016), and that host specific interactions with mutualists and pathogens strongly influence plant population growth (Bennett et al. 2017) and community structure (Bagchi et al. 2014). While such studies have significantly altered past views about the importance of symbiotic interactions during community assembly (Boucher 1982), they have generally taken a phenomenological view with respect to the identity and assembly of the complex microbial communities that underpin emergent effects on host populations and communities. As a result, many questions remain about the capacity of plants to recruit symbiotic microbes in natural systems and how the microbial community assembly process itself influences host plant dynamics.

Understanding how differences in the structure of host-associated microbial communities arise is also important due to the influence these associations exert over ecosystem level processes, such as productivity and nutrient cycling. While microbes are widely acknowledged as direct agents of Earth’s geochemical cycles (Falkowski et al. 2008), there has been significant disagreement about whether microbial community structure *per se* exerts sufficient influence on these same processes, independent of the effect of environmental conditions on microbial communities and physiology (Schimel 1995, Strickland et al. 2009, Maynard et al. 2018). Recent studies have highlighted the impact of mycorrhizal symbioses in particular, with the presence of ectomycorrhizal fungi associated with increases in soil carbon storage (Clemmensen et al. 2013, Averill et al. 2014) and slower nitrogen cycling (Phillips et al. 2013, Corrales et al. 2016, Waring et al. 2016). However, the mechanisms by which plant communities come to be dominated by one symbiosis over another are not well established, and thus it is not known to what extent these are climatically determined (e.g. Houlton et al. 2008, Hedin et al. 2009) or whether they co-exist as potential alternative stable states that result from local biotic feedbacks (Phillips et al. 2013, Corrales et al. 2016, Bennett et al. 2017).

Whether alternative stable states are common in nature, how they arise, and what maintains them is a much-debated topic in ecology (Connell and Sousa 1983, Shurin et al. 2004). In cases where alternative stable states have been documented, positive biotic feedbacks are often invoked: for example, grasses can increase local fire frequency, which in turn promotes their dominance over trees (D’Antonio and Vitousek 1992). However, at least one theoretical model has indicated that positive biotic feedbacks are insufficient to produce stable coexistence of alternative states in the absence of some niche differences and abiotic heterogeneity (Shurin et al. 2004). With respect to symbiotic microbes, on one hand climate appears to have strong effects on the latitudinal distribution of particular symbiotic associations: arbuscular mycorrhizal fungi and nitrogen fixation strategies are most common in tropical forest ecosystems, while ectomycorrhizal symbioses are common in temperate and boreal forests, suggesting important niche differences (Menge et al. 2014, Gei et al. 2018, Steidinger et al. in revision). However, in the tropics large patches of monodominant, ectomycorrhizal forest develop directly adjacent to high diversity, arbuscular mycorrhizal forests under apparently identical initial edaphic conditions (Torti et al. 2001, Henkel 2003, Corrales et al. 2016), suggesting positive biotic feedbacks (Fukami et al. 2017) . Whether alternative stable states arise in soil microbial communities is a key question in plant symbiosis research because it is at the heart of whether or not microbes need to be explicitly accounted for during plant community assembly. If microbial communities simply track climate or the local abiotic environment then their community dynamics can be largely ignored. If, however, functionally distinct microbial communities can arise under identical environmental conditions, then identifying key symbiotic microbes and the limits on their distributions is critical for understanding plant establishment and ecosystem development.

To determine whether stable, alternative soil microbial communities can develop, and whether these alternative communities influence plant establishment and ecosystem function, we conducted a series of field and laboratory experiments in a coastal ecosystem where nearly monodominant stands of plant species coexist across a patchy landscape. At Point Reyes National Seashore (CA, USA), three woody plants species dominate the coastal landscape −*Baccharis pilularis* (BP), which occupies 39% of the study area, *Pinus muricata* (PM; 17%), and *Ceanothus thyrsiflorus* (CT, 18%) (Forrestel et al. 2011). These plants also host distinct combinations of fungal and bacterial root symbioses, either arbuscular mycorrhizal fungi (*B. pilularis, C. thyrsiflorus*), nitrogen fixing actinorhizal bacteria (C. *thyrsiflorus*), or ectomycorrhizal fungi (*P. muricata*). We used historical vegetation data to identify areas in the current landscape that had previously been *B. pilularis* prior to a large fire in 1995. Holding other factors constant (e.g. geology, burn severity, slope), we identified 10 distinct vegetation patches with similar initial environments but representing independent transitions from *B. pilularis* to *P. muricata, B. pilularis* to *C. thyrsiflorus*, or *B. pilularis* to *B. pilularis* (30 total). Sites were also evenly stratified across two geological formations with different properties (sandstone vs. quartz) to compare the relative magnitude of this key abiotic factor. To test whether stable, alternative microbial communities can develop from identical initial conditions, we conducted a DNA based characterization of fungal and bacterial community structure across these sites, as well as measuring a range of ecosystem functions related to carbon and nutrient cycling. Based on the field data, we conducted a factorial greenhouse bioassay experiment where seedlings of the dominant species were grown in each other’s soils, or a mixed soil, to further separate the direct effects of direct plant selection vs. environmental selection on microbial community structure. Finally, we synthesized data from our previous studies to identify the keystone role of particular *P. muricata* fungal mutualists identified as indicator species from community sequencing. Specifically we answer the following questions: (i) given initially identical climate and geology, is a change in plant species sufficient to generate alternative states of microbial community and ecosystem function?; (ii) to what extent are alternative soil microbial communities the result of direct plant effects on microbes versus indirect effects on the soil environment?, and (iii) does the distribution of key microbial taxa also influence transitions between plant community states?

## Materials and Methods

*Study System*. Our study sites are located at Point Reyes National Seashore (PRNS), in Marin County California, USA. The area displays a Mediterranean climate with warm dry summers and cool rainy winters. Average winter low temperatures range between lows of 6-9 °C and highs of 14-17 °C, with summer lows of 10-11 °C and highs of 22-23 °C. Average monthly precipitation ranges from 10.5 to 16 cm during winter and 0.7 to 1 cm during summer. Coastal vegetation at Point Reyes is characterized by a complex matrix of scrub, grasslands and forest. Dominant plant species within each vegetation type, such as Bishop pine (*Pinus muricata* D. Don), Douglas-fir (*Pseudotsuga menziesii* (Mirb.) Franco), Coyote brush (*Baccharis pilularis* DC), and Blue blossom ceanothus (*Ceanothus thyrsiflorus* Eschsch.) tend to form monospecific stands . The vegetation mosaic at PRNS is highly dynamic and strongly influenced by fire (Forrestel et al. 2011), and many of the dominant species have fire adapted recruitment strategies - *P. muricata* has serotinous cones, *C. thyrsiflorus* has a fire activated seed bank, and *B. pilularis* both re-sprouts and produces abundant, small wind dispersed seeds.

### Field Experimental Design

In order to understand whether dominant plant species can create alternative stable states in the microbial community and ecosystem function, we sampled 30 separate stands with a known recent history of vegetative transitions. In 1995, the Vision Fire burned 12,354 acres of lands at PRNS and led to major changes in the composition and spatial distribution of the coastal vegetation matrix at PRNS (Forrestel et al. 2011, Harvey and Holzman 2014). These changes were documented at the meter scale using pre- and post-fire plot data and remote sensing imagery (Forrestel et al. 2011). We use pre- and post-fire vegetation maps to identify independent stands of vegetation representing three pre-fire to post-fire transitions: *Baccharis pilularis* to *Baccharis pilularis*, *Baccharis pilularis* to *Ceanothus thyrsiflorus* and *Baccharis pilularis* to *Pinus muricata*. We randomly selected 10 plots per transition type (30 total) with the following constraints: distance from an established trail less than 300 meters for accessibility; a stand area greater than one acre to limit edge effects; a Normalized Burn Ratio between −150 and 150 to control for major differences in fire intensity. In addition, to examine how differences in the underlying geological substrate influence development of microbial community and ecosystem functions, we stratified our sampling so that 5 of each vegetation transition were sampled from soils derived from Inverness loam (weathered granite) and five soils derived from the Pablo Bayview complex (siliceous shale and sandstone). Location of these soil series were determined from digitization of the USDA Soil Survey of Marin (1985).

### Field Sampling

In each plot we established a sampling grid consisting of three 6m transects. Transects were located a minimum of 5 meters away from each other, although orientation varied somewhat depending on the shape of each patch. Three soil cores (18cm by Ø6cm) were collected at 3m intervals along each transect. Each core was placed in a plastic bag and stored on ice in a cooler until returned to the lab. In the lab, aliquots of soil were frozen at - 80° C for microbial community characterization (see *Molecular Methods*) and the rest used for measuring soil properties. In addition to the nine soil cores, we collected three litter samples (one from each transect) as well as fresh leaves. Litter and leaf tissue were kept at 4° C until further analyses. In addition to soil samples, four resin bags were buried for three months at a depth of approximately 5 cm at each end of the first and the third transect. Two litter bags were also staked on the forest floor and left for three months in each site to measure decomposition rates. One of the two litter bags contained a pure cellulose filter disc (Whatman No. 2) and the other contained 4 grams of dried leaves from the dominant plant species at each plot. Soil, leaf and litter samples were weighed and then dried for 48 hours at 65° C to estimate moisture content. Dried soil and litter samples were ground and total carbon and nitrogen content were measured using on a Carlo-Erba NA 1500 Elemental Analyzers. Nitrate (NO_3_) and Ammonium (NH_4_) from buried resin bags were extracted using 2M KCL and measured using a WestCo SmartChem 20 discrete analyzer. Phosphate-P (PO_4_) was measured on the discrete analyzer after 0.5M HCl extraction from a resin bag incubated with water and 5 g fresh soil for 24 hours (the suspension was shaken at an oscillation rate of 100rpm). All measured chemical variables were averaged to provide a single value for each plot. Establishment of field plots was carried out between November and December of 2014 and litter and resin bags collected in February and March of 2015.

### Soil Bioassay Experiment

In order to decouple the influence of abiotic environment and host species identity on soil microbial community structure, we conducted a greenhouse bioassay using seedlings and field-collected soils from our three dominant plant species (*B. pilularis*, *C. thyrsiflorus, P. muricata*). Seedlings of *P. muricata* and *B. pilularis* were germinated from seeds collected at PRNS. *P. muricata* seeds were germinated using the following steps: seeds were placed in a glass beaker with a 15% v:v dilution of a standard 30% H_2_O_2_ and a drop of Tween. The mixture was placed on an autostirrer for 15-20 minutes. Then the seeds were rinsed with DDI water in a metal strainer, placed in a glass beaker on an autostirrer and allowed to soak overnight (24-48 hours). Finally, the seeds were put between two wet filter paper discs in Petri dishes in a growth chamber (16hr light, held at 21-23°C; day and night). *B. pilularis* seeds were sown directly into wet growing medium consisting of a mix of autoclaved v/v 1/3 vermiculite; 1/3 perlite; 1/3 peat moss. When *P. muricata* and *B. pilularis* seedlings first leaves appeared, they were transferred to pots (Ø 4cm by14cm) filled with an equal mix of live field soil and sterile sand (v/v). Due to a lack of available seeds within the timeframe of the experiment, *C. thirsyflorus* plants were grown from field cuttings collected at PRNS. Cut ends of the *Ceanothus* cuttings were moistened and then dipped in Garden Safe TakeRoot Rooting Hormone and put in autoclaved growing medium (v/v 1/3 vermiculite; 1/3 perlite; 1/3 peat moss). After 3 weeks they were transferred into pots filled with half soil, half sterile sand, as with *P. muricata* and *B. pilularis*.

To see whether microbial communities responded more to the abiotic soil environment or the identity of the dominant plant species, seedlings from each species were grown in factorial combinations of live field soils collected beneath *B. pilularis, C. thyrsiflorus, P. muricata*, or a mix of all three soils. (3 seedling species x 4 soil treatments x 12 replicates = 144). Seedlings were kept in a growth chamber (16hr light, held at 21-23°C; day and night) and watered weekly. After six months seedlings were harvested to characterize microbial community composition and biomass. From each pot we collected 0.25 g of soil and 0.25 g fresh weight roots. The remaining root and shoot tissue from each plant was oven dried and weighed.

### Community metabarcoding

To characterize community composition in field soil, greenhouse soils and roots, we sequenced fungal ITS1 and bacterial 16S rRNA gene amplicons using a multiplexed Illumina library preparation protocol (Smith and Peay 2014). DNA was extracted from field samples using the MoBio PowerSoil RNA kit with DNA elution adapter. DNA was extracted from greenhouse root and soil samples using the MoBio Powersoil kit (MoBio, Carlsbad CA). Field samples were pooled by transect yielding 3 aggregate DNA samples per site.

We amplified the fungal ITS1 region using Illumina primers based on the ITS1F/ITS2 primer pair (forward: 5′-AATGATACGG CGACCACCGA GATCTACACG GCTTGGTCAT TTAGAGGAAG TAA-3′; reverse: 5′-CAAGCAGAAG ACGGCATACG AGAT [INDEX] CGGCTGCGTT CTTCATCGAT GC-3′, where [INDEX] is a sample-specific 12-nt error-correcting Golay barcode). Each reaction contained 0.5–5 μL template DNA, 0.65 U OneTaq HotStart (New England BioLabs), 0.2 μM forward primer, 0.2 μM reverse primer, 0.2 mM dNTPs, and 1 × ThermoPol buffer (New England BioLabs) in a total volume of 25 μL. PCR was performed under the following conditions: initial denaturation at 95°C for 3 minutes, then 35 cycles of denaturation at 95°C for 45 seconds, annealing at 50°C for 45 seconds, and extension at 68°C for 60 seconds, followed by final extension at 68°C for 8 minutes.

For the same samples, we amplified the V4 region of the bacterial 16S rRNA gene using Illumina primers based on the 515F/806R primer pair (forward: 5′-AATGATACGG CGACCACCGA GATCTACACT ATGGTAATTG TGTGCCAGCM GCCGCGGTAA-3′; reverse: 5′-CAAGCAGAAG ACGGCATACG AGAT [INDEX] AGTCAGTCAG CCGGACTACH VGGGTWTCTA AT-3′). To allow additional samples to be multiplexed on the same run, we amplified some samples using a modified forward primer (5′-AATGATACGG CGACCACCGA GATCTACACA CGACATCTAT GGTAATTGTG TGCCAGCMGC CGCGGTAA-3′) containing an 8-base i5 barcode. The reaction mixture was identical to that described above for ITS1. PCR was performed under the following conditions: initial denaturation at 95°C for 3 minutes, then 35 cycles of denaturation at 95°C for 45 seconds, annealing at 51°C for 60 seconds, and extension at 68°C for 90 seconds, followed by final extension at 68°C for 5 minutes.

PCR products were checked using agarose gel electrophoresis. For each batch of PCR reactions, we visually inspected 16S and ITS1 amplicon bands and pooled product volumes according to intensity (1 μL for samples with the strongest bands, up to 3 μL for those with weak bands). To remove unincorporated primers and primer dimers, we cleaned pooled DNA from each batch using Sera-Mag SpeedBeads (Fisher Scientific #65152105050250) according to the size-selection protocol in Rohland and Reich (2012). Finally, we combined the pooled DNA into three mixed ITS1/16S libraries with final DNA concentrations of 4 nM. We quantified DNA concentrations with the Qubit dsDNA HS Assay Kit (Life Technologies # Q33230) and calculated volumes needed to ensure equimolar amounts of DNA were included from each reaction. We adjusted volumes so that 0.5 times more amplicons per sample were included for ITS1 amplifications than for 16S amplifications due to previously observed bias in sequencing efficiency. The three mixed ITS1-16S libraries were sequenced separately on the Illumina MiSeq platform (paired-end reads, 2×300 cycles) at the Stanford Functional Genomics Facility. We used custom sequencing primers for the R1 read (ITS1: 5′-TTGGTCATTT AGAGGAAGTA AAAGTCGTAA CAAGGTTTCC-3′; 16S: 5′-TATGGTAATT GTGTGCCAGC MGCCGCGGTA A-3′), R2 read (ITS1: 5′-CGTTCTTCAT CGATGCVAGA RCCAAGAGAT-3′; 16S: 5′-AGTCAGTCAG CCGGACTACH VGGGTWTCTA AT-3′), and P7 index read (ITS1: 5′-TCTCGCATCG ATGAAGAACG CAGCCG-3′; 16S: 5′-ATTAGAWACC CBDGTAGTCC GGCTGACTGA CT-3′). Additionally, we sequenced the P5 index using the standard Illumina i5 sequencing primer (only used to identify samples amplified using the alternate forward 16S primer).

### Metabarcoding analysis

Reads were demultiplexed and assigned to samples using the default Illumina BaseSpace software. After separating ITS1 and 16S samples, we processed data from each marker and study separately. Using VSEARCH (Rognes et al. 2016), we trimmed reads after a quality score of 5 or below, requiring that at least 150 bases remained and there was at most 1 expected error. Read pairs with at least 16 bp of overlap where at most 10% of overlapping bases were different were merged if the resulting sequence was at least 75 bp. Remaining adaptors were trimmed with cutadapt (Martin 2011). Finally, using VSEARCH, we clustered sequences into 97% sequence identity OTUs. Taxonomic identity was assigned for a representative sequence for each OTU using the Ribosomal Database Project naïve Bayesian classifier (Wang et al. 2007) with a bootstrap confidence of 50%. Classification for 16S used the RDP 16S train set 16 and the fungal ITS OTUs the Warcup train set 1 (Deshpande et al. 2016). Taxonomy, OTU table, and sample information were combined into a single R object for downstream manipulation using Phyloseq (McMurdie and Holmes 2013). For all datasets, potential contaminant OTUs were identified statistically based on prevalence in negative control samples (both PCR and extraction controls) for ITS and 16S libraries using the R package decontam (Davis et al. 2017) based on OTU prevalence and a threshold of 0.1. For the field samples this resulted in the elimination of 125 of ITS and 7 16S OTUs. For the bioassay samples this resulted in elimination of 49 of 11,852 ITS and 149 of 13,1488 16S OTUs. For the field study we next removed samples with <5000 sequences and all OTUs represented by fewer than 100 sequences to focus the dataset on the core microbial community. Samples were then merged within each plot and rarefied to an even depth of 65,000 and 5,000 sequences for ITS and 48,000 and 2,500 sequences for 16S for field and bioassay experiment, respectively. All OTUs identified as Chloroplast or Mitochondria were removed from the 16S datasets prior to analysis.

### Statistical analysis

We used analysis of variance (ANOVA) to test whether dominant vegetation, geological substrate, or their interaction influenced the ecosystem properties we measured. Given the differences we observed, we also used litter weight per area to extrapolate our measurements to the vegetation extent at PRNS (Forrestel et al. 2011).

To analyze field microbial community composition across vegetation stands we used two approaches. First, to test whether vegetation type and geological substrate affected fungal and bacterial composition we used permutational multivariate ANOVA (Anderson 2001) based on Bray Curtis dissimilarities and implemented in the R package vegan (Oksanen et al. 2008). Second, to test which environmental variables were most strongly correlated with microbial community structure we used generalized dissimilarity modeling (GDM), which uses iSplines to model nonlinearity in community change and identify the portion of environmental gradients with the most rapid community change (Manion et al. 2018). We identified taxa associated with different plant species in both the field and greenhouse study using differential expression analysis based on the negative binomial distribution as implemented in the DESeq2 package (Love et al. 2014) using an adjusted p-value of <0.05. Overlap of the most common taxa (>1% of sequences) and indicator species (p < 0.05) across field and greenhouse components of the study were visualized using circular diagrams (Gu et al. 2014). Although we ended up with low replication for sequencing greenhouse *C. thyrsiflorus* seedling treatments, repeating community analyses without *C. thyrsiflorus* seedlings did not qualitatively change our results (data not shown) so we present results from the full analysis.

Because DESeq2 indicated minimal overlap between indicator taxa in mature vegetation and seedlings, we wanted to further examine the potential ecological importance of these seedling associated taxa. While detailed ecological data is absent for most microbial species, because of our long history of work at PRNS, we were able to synthesize data for a small group of ectomycorrhizal fungi, the Suillineae, consisting of two genera (*Rhizopogon* & *Suillus*) and seven species at PRNS. First, we summarized their relative abundance in soil sequence datasets from *P. muricata* seedlings (this study), 20 year forest (this study), and mature (>50 year) forest (Talbot et al. 2014). Next, we examined their importance as highly dispersive mutualists by calculating the fraction of *P. muricata* seedlings recruiting in *B. pilularis* soils that would have gone uncolonized by ectomycorrhizal fungi in the absence of the Suillineae, based on data from Smith et al. (2018). Because we know that the Suillineae are generally poor competitors (Lilleskov and Bruns 2003, Kennedy et al. 2011, Smith et al. 2018), seedlings colonized only by Suillineae would not have otherwise been colonized by other species of ectomycorrhizal fungi. Because the design of the Smith et al. (2018) study was spatially explicit, we are able to calculate how the fraction of colonized seedlings changes for pine seedlings establishing in *B. pilularis* vegetation as their distance away from established *P. muricata* vegetation increases. Finally, to examine the importance of these fungi during the early stages of plant competition, we used data from (Peay 2018) to calculate the biomass of *P. muricata* and *B. pilularis* seedlings when competing against each other in natural *B. pilularis* soils and in *B. pilularis* soils amended with spores from a native species of Suillineae (*Suillus pungens*). These soils are derived from a location distant to established *P. muricata* and naturally low in ectomycorrhizal inoculum, allowing us to ask how plant competitive dynamics might change at PRNS in the absence of the Suillineae seedling colonization.

## Results & Discussion

In the >20 years since the 1995 Vision Fire, sites that transitioned from vegetation dominated by *Baccharis pilularis* to dominance by *Pinus muricata* or *Ceanothus thyrsiflorus* developed considerable differences in a range of ecosystem properties. This was evidenced by significant effects of the vegetation transition for 10 of the 15 variables we measured, including the speed of decomposition, the quality and quantity of litter, and the concentration of nitrogen and carbon in soils (*Table S1*, **Fig 1**), with vegetation type explaining on average 31% of variation (range = 5 - 69%). By contrast, difference in the underlying geological substrate, or interaction between vegetation and geology, were significant in only four cases (*Table S1*, **Fig S2, S3**), explaining an average of only 6% of variation (range = 0 – 32%).

**Figure 1:**
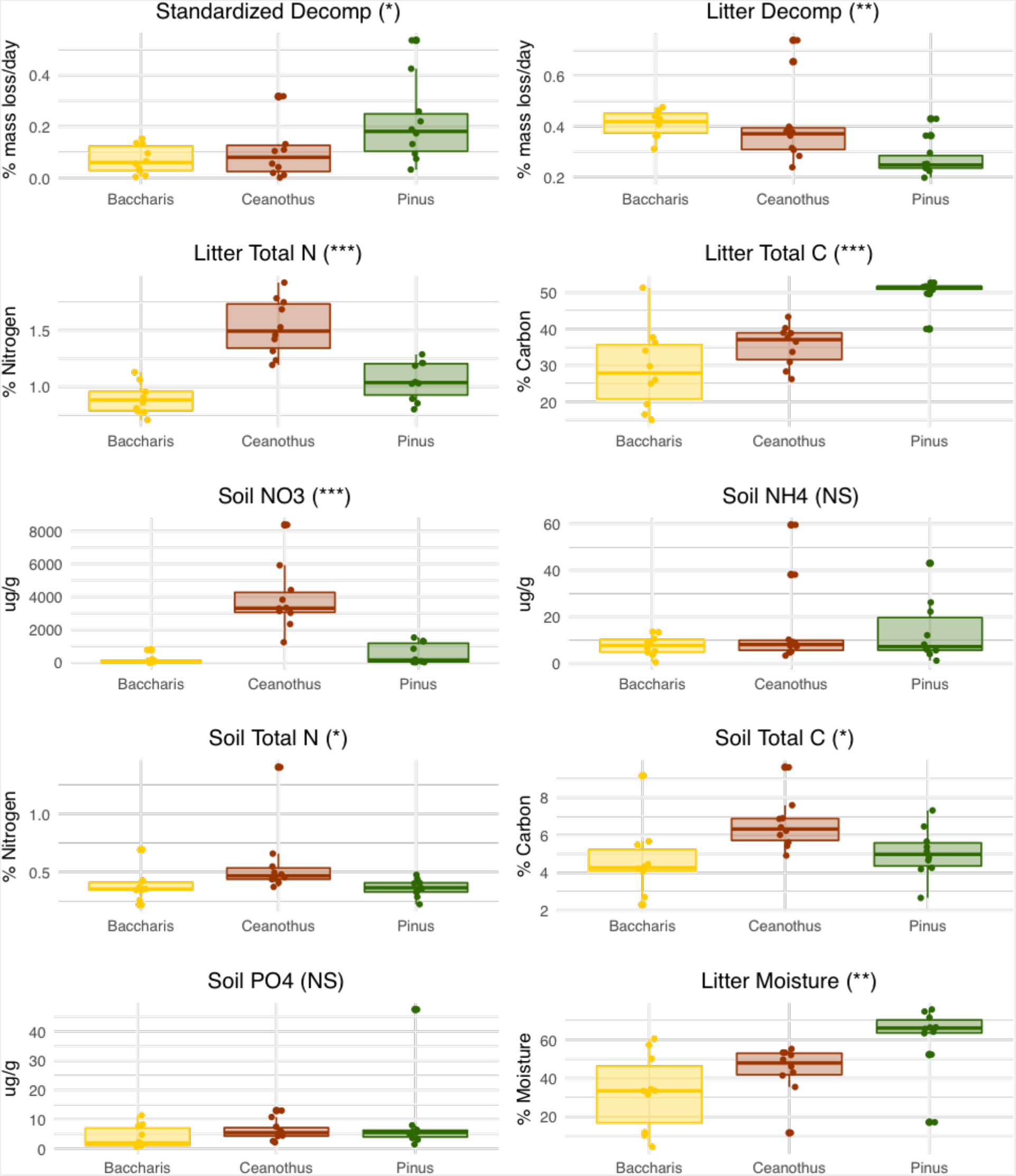
Variation in ecosystem properties after transition between dominant plant species. Bars show mean +/-1 standard error for 10 variables measured under 30 stands that transitioned from *B. pilularis* to either *B. pilularis, C. thyrsiflorus*, or *P. muricata*. P-values are indicated as . < 0.1, * < 0.05, ** < 0.01, and *** < 0.001.

While abiotic variables have often been thought of as the primary constraint on key biological processes, such as productivity or decomposition (Lieth and Whittaker 1975, Schimel 1995, Adair et al. 2008), an emerging body of literature suggests that differences in species abundance or composition may affect these key biological processes to an equal or greater extent (Cornwell et al. 2008, Schnitzer et al. 2011, Allison et al. 2013, Bradford et al. 2014, Terrer et al. 2016). For example, a recent global change field experiment found that microbial community structure explained nearly 3x as much variation as abiotic field conditions when predicting wood decomposition rates (Maynard et al. 2018). Our field data indicate that biotic controls over ecosystem properties are the product of trait differences between both primary producers and the activity of their associated heterotrophic soil microbes. For example, litter quality (C and N content) varied substantially across the three dominant plant species. This likely directly impacted decomposition rates, as evidenced by the slowest home litter decomposition rates beneath pines, which had the highest litter C:N ratio, but also indirectly by altering the physiological potential of the microbial community, as we found that the standard substrate (pure cellulose) by contrast decayed much more rapidly beneath pines (**Fig 1**). Similarly, the ability of *Ceanothus* to host N-fixing bacteria clearly elevated both tissue and soil N concentrations. Such differences are biologically significant, translating to an estimated increase to date of 4,362 metric tons of C and 167 tons of N in litter across the study area as a result of conversion from *B. pilularis* to *P. muricata* and *C. thyrsiflorus* after the 1995 fire. In aggregate, these differences show that clearly distinct states of the soil ecosystem are initiated through the establishment of dominant plant species and can co-occur contemporaneously across a landscape even under identical starting conditions.

Changes in soil properties and processes after transition from *B. pilularis* to either *P. muricata* or *C. thyrsiflorus* were matched by large changes in the composition of fungi and bacteria. Differences in community composition were more pronounced for fungal communities, which were distinct across all vegetation types (P < 0.001, R^2^ = 0.18), compared with bacteria, which showed overall weaker effects of vegetation type (P = 0.012, R^2^=0.12) and greater similarity between *P. muricata* and *C. thyrsiflorus* (**Fig 2 a,b**; *Table 1, Table 2*). Bacteria also showed a significant effect of geological substrate (P =0.023, R^2^ = 0.07) that was not evident for fungi (P =0.445, R^2^ = 0.03). Other studies have also found that fungal communities are more strongly influenced by the dominant vegetation than bacterial communities are, consistent with the possibility that fungi have greater host or substrate specificity than bacteria (Urbanová et al. 2015). Changes in fungal community composition were also well explained by stand level environmental variables, such as litter chemistry and soil moisture (GDM R^2^ = 0.30; **Fig 2c**; *Table S2*) that were correlated with changes in vegetation. Bacterial community change was best predicted by pH and litter mass, environmental factors that showed more complex, interactive variation related to both vegetation type and geology (GDM R^2^ = 0.21; **Fig. 2d**; **Fig S3**; *Table S3*). GDM partial plots show that microbial communities exhibit a range of response functions to environmental variation. For instance, community change was relatively continuous with respect to litter C:N ratio (fungi) and pH (bacteria), but showed threshold effects with relatively small initial increases in soil moisture (fungi) and litter depth (bacteria). Because GDM is able to identify where along a gradient community change occurs it can provide additional insight compared with standard distance-based methods of community analysis. While relatively few microbial community studies (Glassman et al. 2017) have applied this approach, the existence of multiple response functions may help explain the high β-diversity of soil microbial communities.

**Figure 2.**
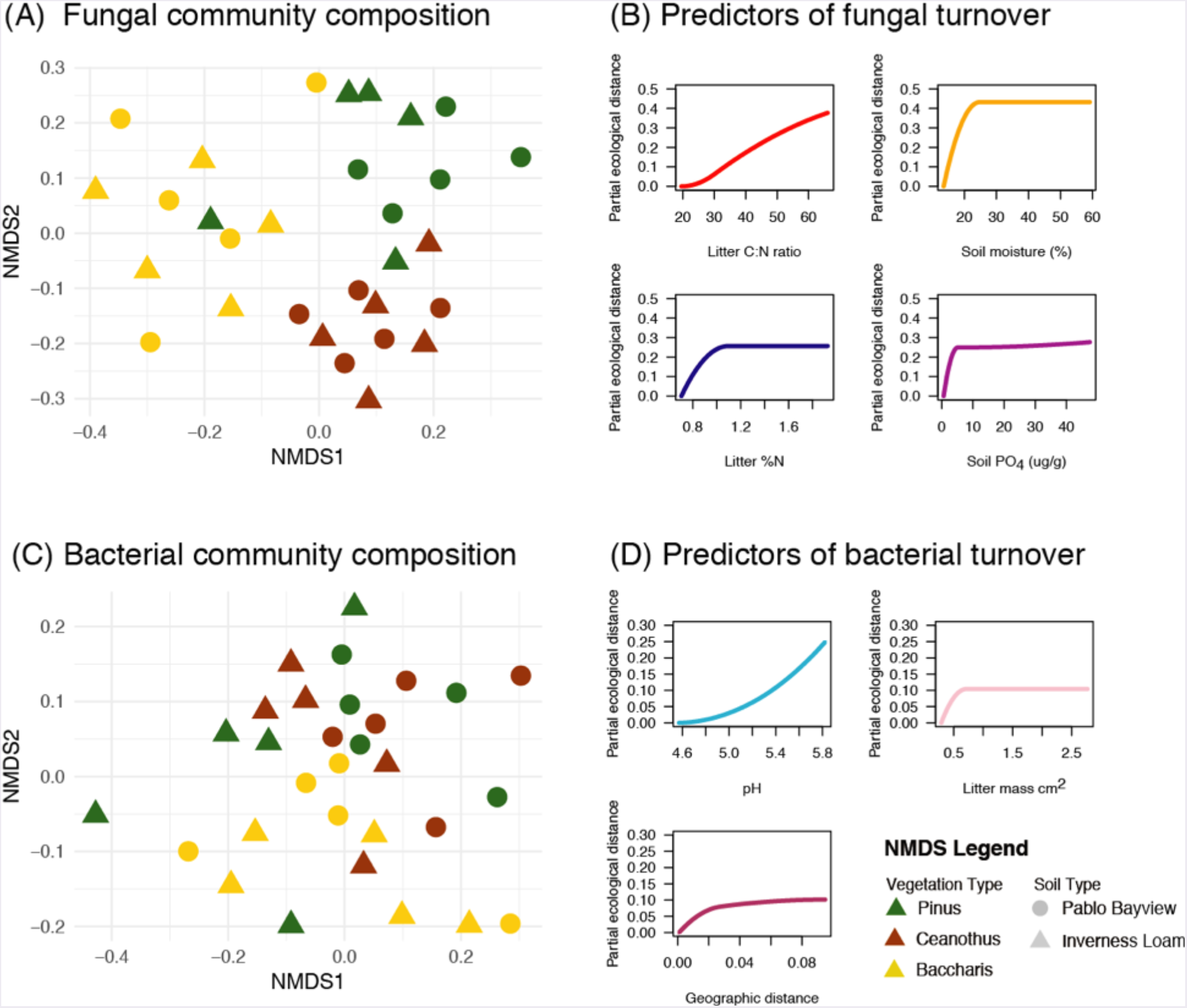
Variation in microbial community composition after transition between dominant plant species. Ordination of (a) fungal and (c) bacterial communities sampled from field soils at Point Reyes National Seashore. Each point represents an independent patch that transitioned from *B. pilularis* to either *B. pilularis*, *C. thyrsiflorus*, or *P. muricata* following the 1995 Vision Fire. Generalized dissimilarity model shows significant environmental predictors of fungal (b) and bacterial (c) community change. The magnitude of change along the y-axis indicates the relative importance of variables, while slope indicates portions of the environmental gradient where community change is most rapid.

**Table 1.**
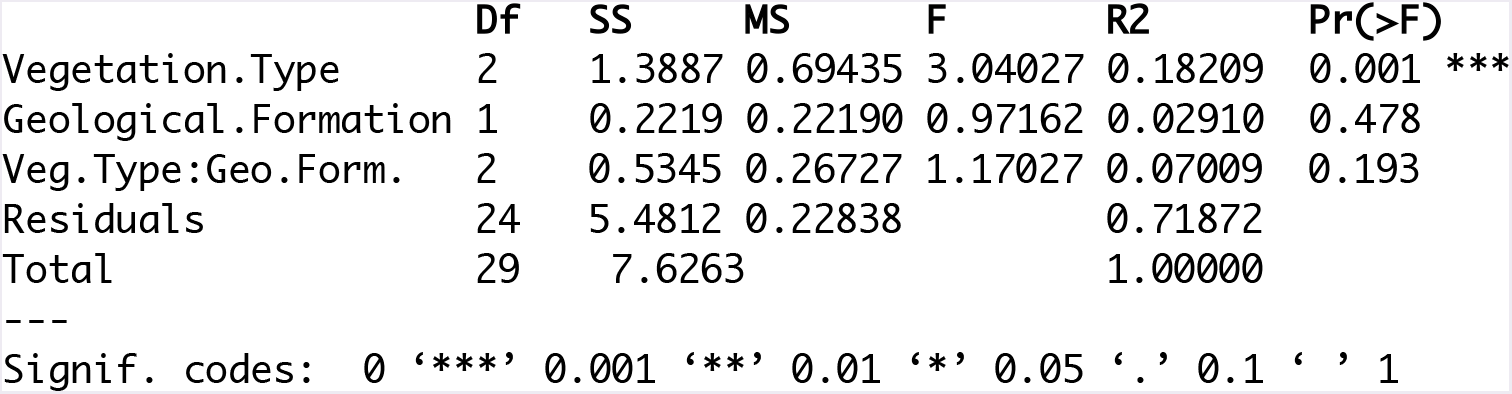
Field fungal communities are strongly influenced by vegetation type. Summary table from permutational multivariate ANOVA of fungal communities from field plots at Point Reyes National Seashore. Results show the effect of three vegetation types (B. *pilularis, C. thyrsiflorus, P. muricata*), two geological formations (Inverness Loam, Pablo Bayview Complex), and their interaction.

**Table 2.** Field bacterial communities are influenced by vegetation type. Summary table from permutational multivariate ANOVA of bacterial communities from field plots at Point Reyes National Seashore. Results show the effect of three vegetation types (B. *pilularis, C. thyrsiflorus, P. muricata*), two geological formations (Inverness Loam, Pablo Bayview Complex), and their interaction.

|  | Df | SS | MS | F.Model | R2 | Pr(>F) |
| --- | --- | --- | --- | --- | --- | --- |
| Veg.Type | 2 | 0.28011 | 0.140057 | 1.9678 | 0.12249 | 0.021 * |
| Geological.Formation | 1 | 0.13060 | 0.130601 | 1.8349 | 0.05711 | 0.062 . |
| Veg.Type:Geo.Form. | 2 | 0.16782 | 0.083909 | 1.1789 | 0.07339 | 0.247 |
| Residuals | 24 | 1.70822 | 0.071176 |  | 0.74701 |  |
| Total | 29 | 2.28676 |  |  | 1.00000 |  |

Signif. codes: 0 '\*\*\*' 0.001 '\*\*' 0.01 '\*' 0.05 '.' 0.1 ' ' 1

Plant and microbial community structure often co-vary across environmental gradients (Peay et al. 2010, Peay et al. 2013, Mueller et al. 2014, Waring et al. 2016). While in many cases this pattern may be driven by independent responses of plants and microbes to external drivers (i.e. climate, geology), plants themselves may also have strong effects local soil conditions (Hooper and Vitousek 1997, Waring et al. 2015). Because there was no pre-existing environmental gradient in this system, given that all patches started as *B. pilularis* and were stratified evenly across the two different geological substrates, the changes in microbial community that we observed are ultimately attributable to changes in the plant community. However, the large changes in the soil abiotic environment associated with plant litter quality make it difficult to assign relative importance between two forms of causality based on field data alone, i.e., does plant modification of the bulk soil environment indirectly select different microbial communities, or are microbial communities directly selected by more specialized symbiotrophic interactions? Previous common garden or bioassay experiments that attempt to control for plant effects on the soil abiotic environment have shown mixed results. For example, reciprocal transplants that break up plant species-environment correlations across nutrient gradients have shown weak effects of plant species identity *per se* on the microbial community (Peay et al. 2015, Erlandson et al. 2018). By contrast, Gehring et al. (2017) showed that plant genotypes with different traits grown in a common environment still selected functionally different ectomycorrhizal fungi. Disentangling between these two types of causality in microbial community assembly has important implications for whether or how microbial communities can be manipulated to manage ecosystem processes, such as carbon sequestration or sustainable agriculture (Busby et al. 2017, Toju et al. 2018).

Put in this perspective, our greenhouse reciprocal transplant bioassay shows experimentally that soil microbial communities are shaped extensively through symbiotic interactions in the plant root system that may eventually extend to transform the entire soil community. After 6 months of growth in live field soil, seedling species exerted significant control over both bacterial and fungal communities. As a result, microbial communities assembled under either *B. pilularis, C. thyrsiflorus*, or *P. muricata* tended to be self-similar regardless of the starting soil inoculum (**Fig 3**, *Tables S4,5*). However, for both bacteria and fungi the strength of plant control varied depending on whether we examined root or soil components of each replicate (*Host Species x Inoculum x Substrate* P < 0.05; *Tables S4, S5*), with root microbial communities most strongly influenced by plant species identity, and bulk soil communities most strongly influenced by inoculum source (*Tables 3-6*). This effect was most pronounced for bacteria, where plant species identity explained 22% of community variation in roots but only 4% in soils. Conversely, inoculum source explained 13% of community variance in roots and 27% in soils. Interestingly, root communities that developed in mixed inoculum (equal parts of soil from all three species) ordinated with plant species growing in their home soils, strongly suggesting co-adaptation of plants with their home soil microbes. Our results are consistent with accumulating evidence that plant species influence considerable control over microbial communities within the rhizosphere, but relatively weak direct control within the bulk soil (Hartmann et al. 2008, Mendes et al. 2014, Nuccio et al. 2016). However, by controlling for soil environment in our experiment and juxtaposing seedlings (6 months) and field communities (>20 years), it is clear that over time root associated microbial communities may extend their influence out to contribute to the broad differences in soil microbial communities seen in soils under different vegetation types (Urbanová et al. 2015). This effect is more evident for fungi, whereas bacteria were not as strongly differentiated in field soils (**Fig 2**) despite strong effects in seedling roots (**Fig 3**, *Table 5*). Because mycorrhizal fungi both colonize roots and grow into the soil to forage for nutrients, their spatial influence extends beyond the root zone. As a result, it is likely that biotic interactions with mycorrhizal fungi play a foundational role in structuring the rest of the soil microbial community.

**Figure 3.**
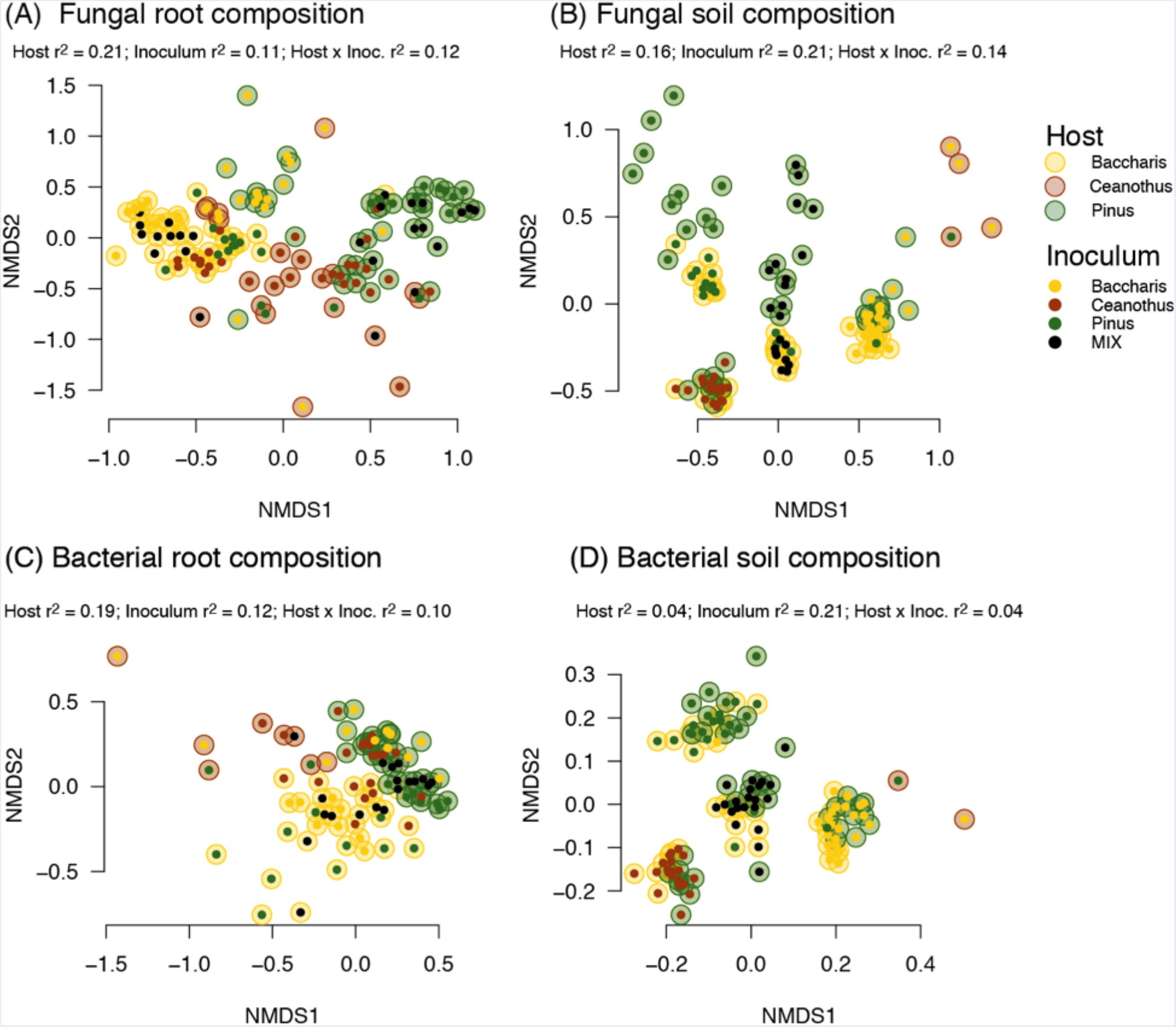
. Microbial communities are built from the inside out. Effects of seedling species identity and inoculum source on the composition of fungi (A,B) and bacteria (C,D) on plant roots (A,C) and soil (B,D) in a greenhouse experiment. Colors indicate plant species: *B. pilularis* (BAPI, yellow), *C. thyrsiflorus* (CETH, red), and *P. muricata* (PIMU, green). Outer circle indicates the identity of seedling species grown in each pot, while the inner circle indicates identity of the plant species from which the soil inoculum was collected beneath. Proportion of variance explained for all significant PERMANOVA terms is given beneath the plot title.

**Table 3:**
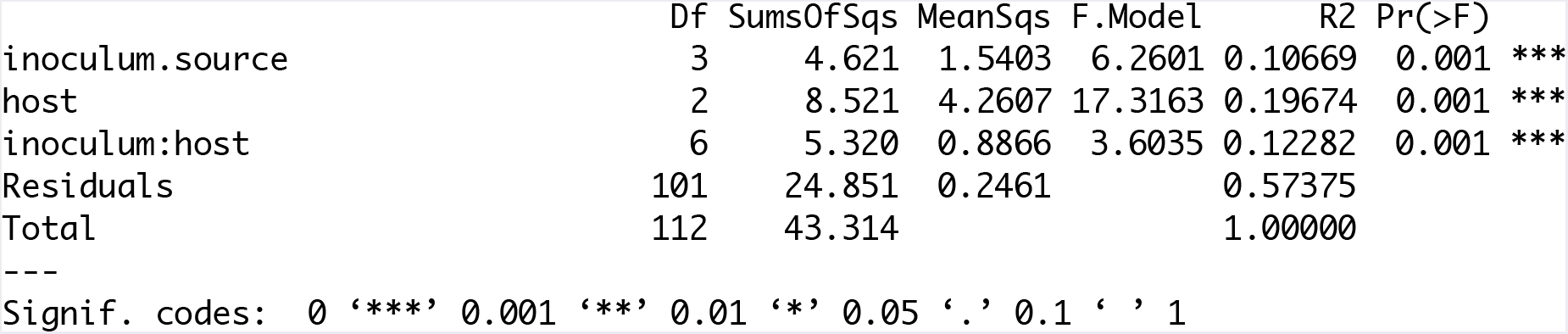
Root fungal community assembly is most influenced by host identity. Summary table from permutational multivariate ANOVA of fungal communities from bioassay seedlings of *B. pilularis, C. thyrsiflorus, P. muricata* (host) grown in live field soils taken from beneath each host (inoculum.source).

**Table 4:**
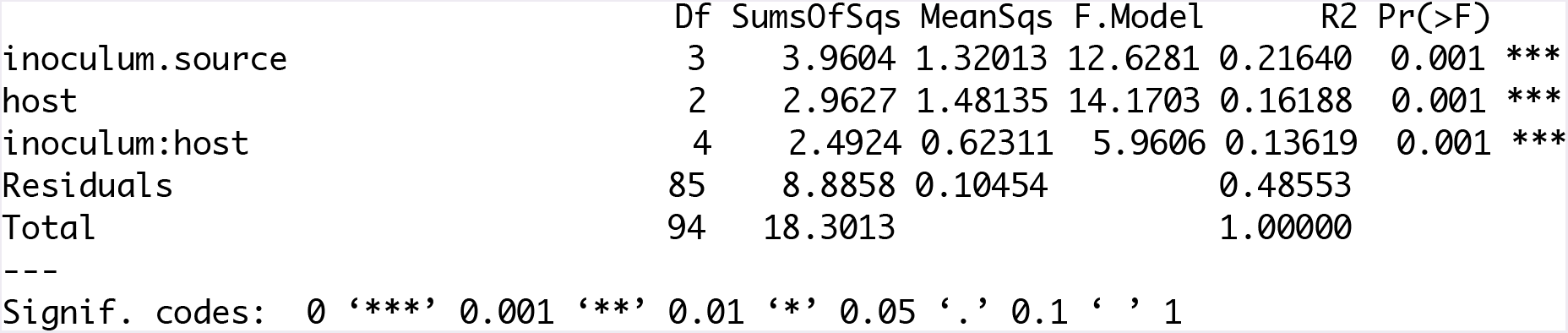
Soil fungal community assembly is most influenced by inoculum source. Summary table from permutational multivariate ANOVA of fungal communities from bioassay seedlings of *B. pilularis, C. thyrsiflorus, P. muricata* (host) grown in live field soils taken from beneath each host (inoculum.source).

**Table 5:** Root bacterial community assembly is most influenced by host identity. Summary table from permutational multivariate ANOVA of bacterial communities from bioassay seedlings of *B. pilularis, C. thyrsiflorus, P. muricata* (host) grown in live field soils taken from beneath each host (inoculum.source).

|  | Df | SumsOfSqs | MeanSqs | F.Model | R2 | Pr(>F) |  |
| --- | --- | --- | --- | --- | --- | --- | --- |
| inoculum.source | 3 | 1.6104 | 0.53681 | 4.8837 | 0.13475 | 0.001 | *** |
| host | 2 | 2.4487 | 1.22434 | 11.1386 | 0.20490 | 0.001 | *** |
| inoculum.source:host | 5 | 1.1866 | 0.23733 | 2.1591 | 0.09929 | 0.001 | *** |
| Residuals | 61 | 6.7050 | 0.10992 |  | 0.56105 |  |  |
| Total | 71 | 11.9508 |  |  | 1.00000 |  |  |
| --- |  |  |  |  |  |  |  |
| Signif. codes: 0 '***' 0.001 '**' 0.01 '*' 0.05 '.' 0.1 ' ' 1 |  |  |  |  |  |  |  |

**Table 6:**
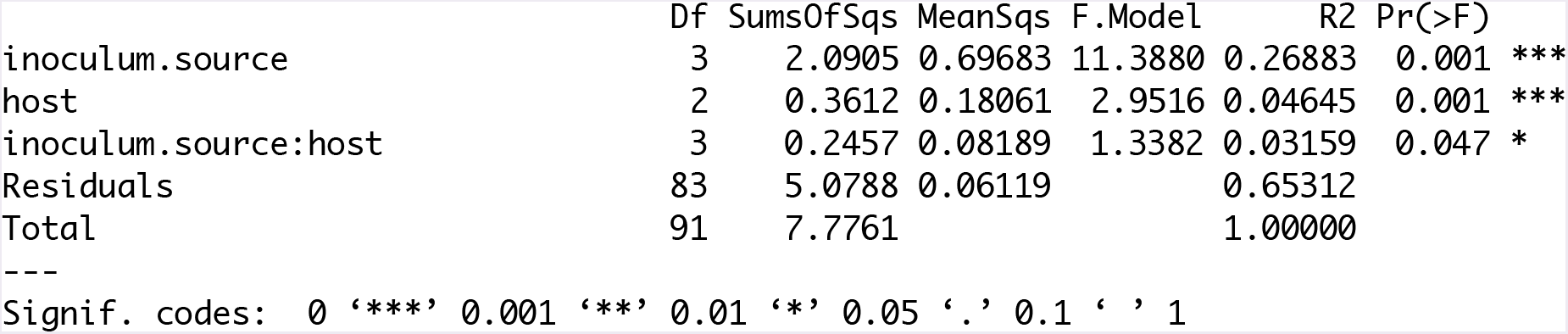
Soil bacterial community assembly is most influenced by inoculum source. Summary table from permutational multivariate ANOVA of bacterial communities from bioassay seedlings of *B. pilularis, C. thyrsiflorus*, *P. muricata* (host) grown in live field soils taken from beneath each host (inoculum.source).

Inspection of the taxa responsible for differentiating plant associated microbial communities, both in field soils and seedling bioassays, demonstrates that particular symbiotrophic taxa may act as keystone species during ecosystem assembly. Although the definition of a keystone species has been debated (Cottee-Jones and Whittaker 2012), keystone species are often defined as species which have a strong trophic impact relative to their biomass (Paine 1969, Power and Mills 1995). There has been growing interest in the identification of keystone microbes using community network properties (Banerjee et al. 2018, Herren and McMahon 2018). While such network-based analyses are critical for rapid identification of keystone taxa in complex communities, validating the ecological impact of putative microbial keystones is also important to put their effects into the same context as other classic macrobial keystone taxa. Our data provides direct evidence that some symbiotrophic microbes, and in particular mycorrhizal fungi, may act as classic keystone species by mediating shifts in the dominant vegetation that have cascading effects on microbial communities and ecosystem functions. First, in our study many of the taxa which significantly differentiated fungal communities in the field and bioassays were mycorrhizal. For example, mycorrhizal fungi made up 20% (12 of 58) and 18% (12 of 67) of taxa identified through differential expression to differentiate *P. muricata* field soils from *B. pilularis* and *C. thyrsiflorus*, and consistently exhibited the largest log fold changes (*SupplementalFiles*). While symbiotrophic fungi can make up a significant fraction of sequence data (26% across our field dataset), individual taxa are generally rare with the most common species, an ectomycorrhizal fungus named *Tomentella sublilacina*, accounting for only 2% of all fungal sequences. Total fungal biomass in soils is also ~40 times lower than that of plants (Bar-On et al. 2018). Second, we observe minimal overlap in community composition between field soils and seedling bioassays (**Fig 4, 5, S6-9**), both in terms of abundance and their identification as differentially abundant taxa: only two significantly enriched taxa were shared between *P. muricata* field soils and *P. muricata* seedlings, no shared taxa discriminated *B. pilularis* field soils and *B. pilularis* seedlings. For example, *Rhizopogon*, a member of the Suillineae that was common on *P. muricata* seedlings invading coastal scrub immediately after the fires at PRNS (Horton et al. 1998), was rare in our field soils but highly common in greenhouse bioassays. The only exception to this pattern was bacterial *C. thyrsiflorus* associated communities, which showed high overlap with *C. thyrsiflorus* seedlings in the greenhouse (**Fig S9**). Thus, these symbiotrophic taxa are generally low in abundance but play a significant role in distinguishing soil microbial communities.

**Figure 4:**
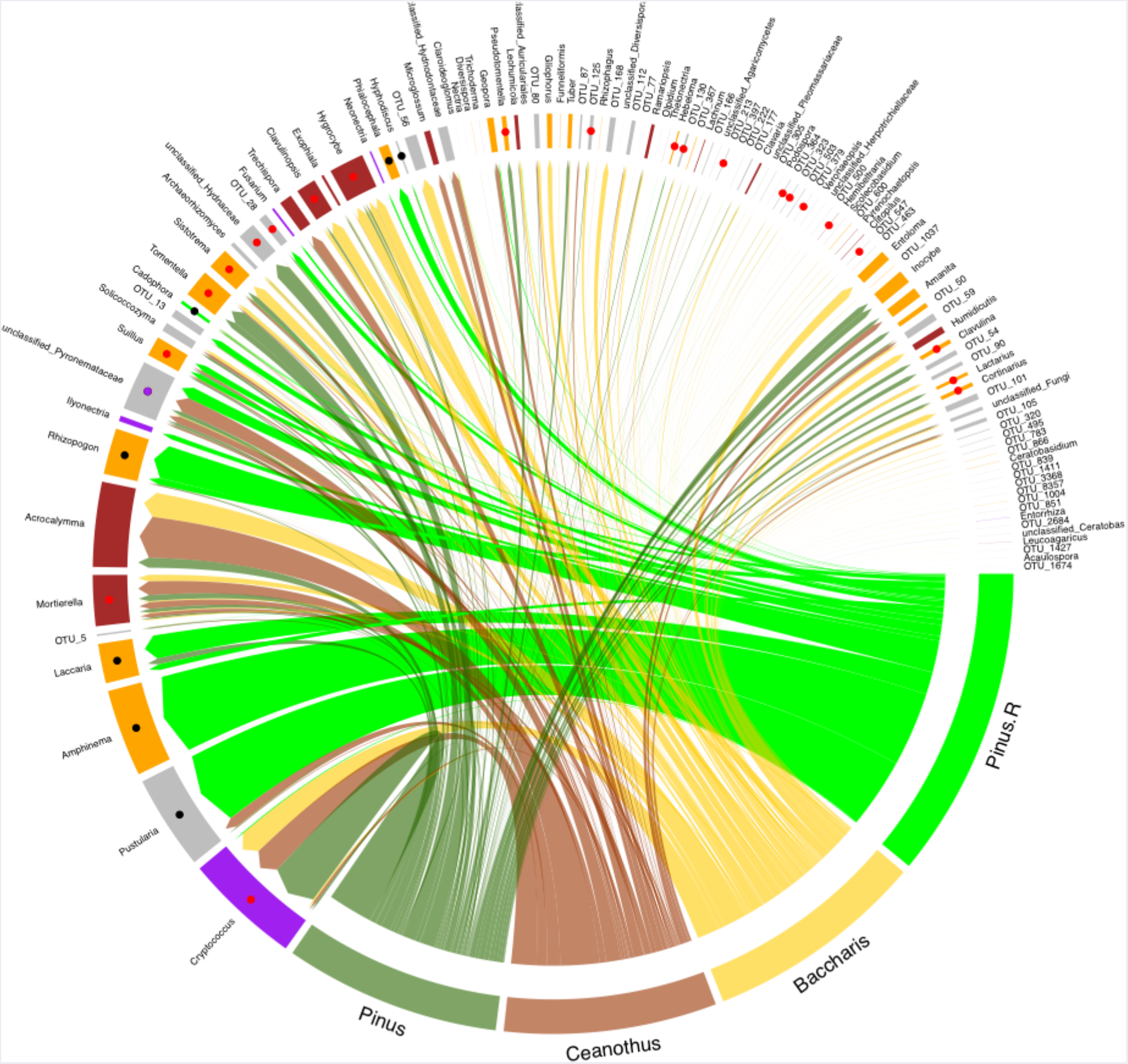
Minimal overlap between fungal communities found on seedling roots and in field soils. Circular diagram shows taxonomic overlap between field soil communities under *P. muricata* (Pinus; light green), *C. thyrsiflorus* (Ceanothus; light red), *B. pilularis* (Baccharis; light yellow) and on *P. muricata* seedling roots (Pinus.R; dark green) in a greenhouse bioassay. Common (>1% sequence abundance) or indicator OTUs are plotted and track color indicates trophic guild: mycorrhizal (orange), saprotrophic (maroon), pathogens (purple), endophytes (green), or unknown (grey). Points show taxa that positively differentiate (adjusted P < 0.05) *P. muricata* field soils from other field soils (red points), positively differentiate *P. muricata* seedling roots from other seedling roots (black points), or differentiate both (purple). Width of track and arrows are proportional to the number of sequences for each OTU.

**Figure 5:**
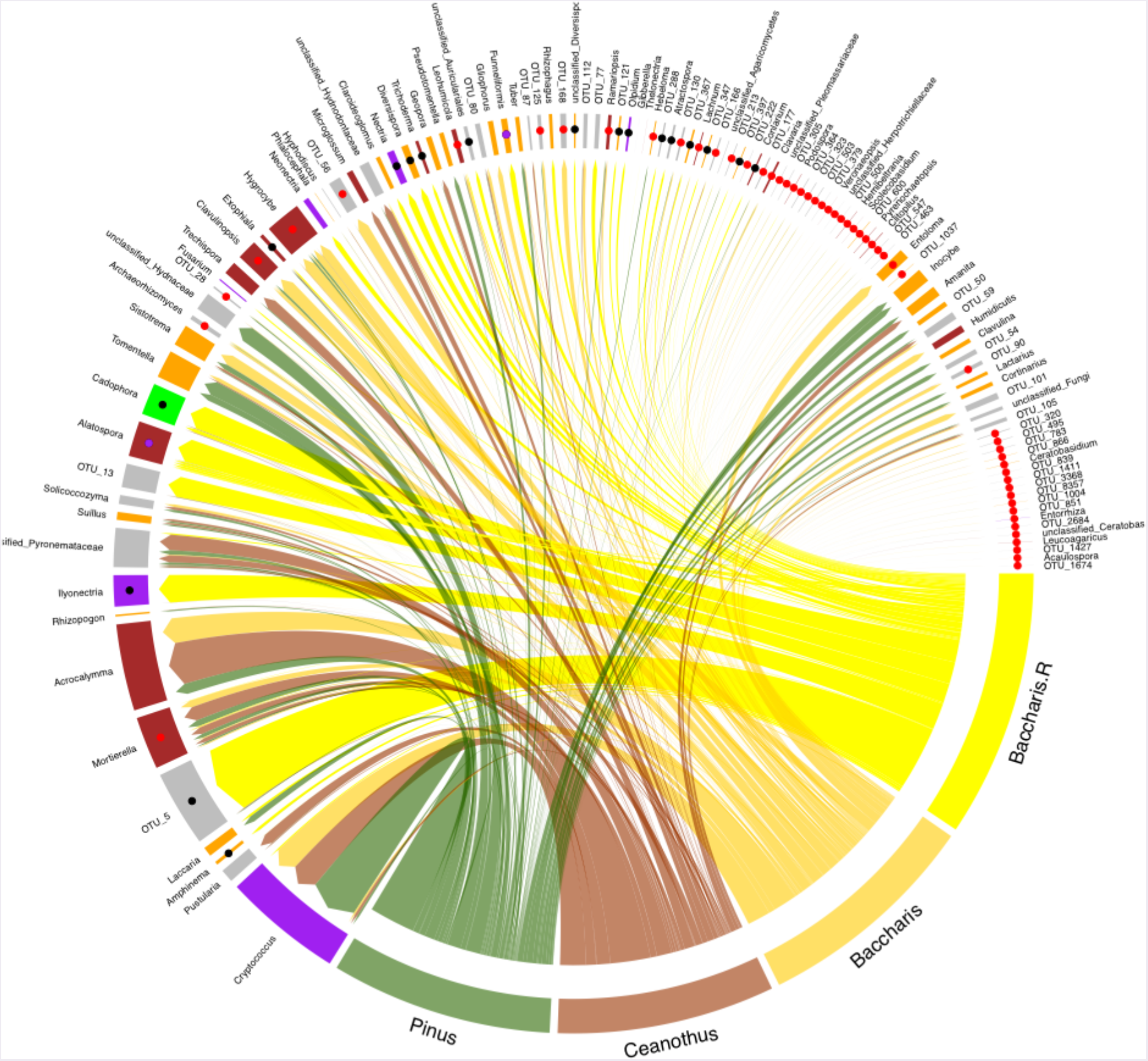
Minimal overlap between fungal communities found on seedling roots and in field soils. Circular diagram shows taxonomic overlap between field soil communities under *P. muricata* (Pinus; light green), *<u>C. thyrsiflorus</u>* (Ceanothus; light red), *B. pilularis* (Baccharis; light yellow) and on *B. pilularis* seedling roots (Baccharis.R; dark yellow) in a greenhouse bioassay. Common (>1% sequence abundance) or indicator OTUs are plotted and track color indicates trophic guild: mycorrhizal (orange), saprotrophic (maroon), pathogens (purple), endophytes (green), or unknown (grey). Points show taxa that positively differentiate (adjusted P < 0.05) *B. pilularis* field soils from other field soils (red points), differentiate *B. pilularis* seedling roots from other seedling roots (black points), or differentiate both (purple). Width of tracks and arrows are proportional to the number of sequences for each OTU.

Given the preponderance of rare taxa in microbial communities, understanding their ecology and impact is a major knowledge gap (Banerjee et al. 2018). Analysis of community change indicates that many taxa are conditionally rare, and that their dynamics contribute disproportionately to community diversity (Shade et al. 2014). Yet for many of these taxa insufficient evidence exists to explain their ecology or importance. We synthesized some of our >10 years of work on ectomycorrhizal fungal communities at PRNS to illustrate the ecology and impact of these conditionally rare taxa. First, many of these conditionally rare microbes are likely ruderal, with strong dispersal and weak competitive abilities. Focusing on a single ectomycorrhizal group for which we have extensive data and were common on *P. muricata* bioassay seedlings, the Suillineae, sequence data shows that while they are common on seedlings in disturbed settings (**Fig 6a**), they are generally rare both in mature pine forests at PRNS and across North America. Second, they are dispersal specialists and, during pine invasion of *B. pilularis* coastal scrub, increasingly dominate pine seedling root communities with increasing distance from established pines (**Fig 6b**). For example, in the absence of the Suillineae we estimate that ectomycorrhizal colonization of pine seedlings establishing in *B. pilularis* vegetation would drop by 60% (from 100% to 40% of seedlings colonized) over only 10 meters distance from established pines. The Suillineae are strict specialists on the Pinaceae and invading pine seedlings may be effectively colonized by aerial dispersal of the Suillineae over 100s to 1000s of meters (Peay et al. 2012, Hayward et al. 2015). By contrast, ectomycorrhizal colonization of oak seedlings establishing in arbuscular mycorrhizal grasslands has been observed to drop off rapidly over a few meters away from established oaks (Dickie and Reich 2005), illustrating the potential impact on seedling colonization dynamics of this group of dispersal specialists. Finally, the ability of these fungi to be present during plant invasion has important consequences for vegetation dynamics. When the presence of Suillineae in *B. pilularis* field soils are experimentally manipulated, pine seedlings are at a competitive disadvantage to *B. pilularis* when they are absent, and competitively dominant when they are present (**Fig 6c**). Given their importance in seedling establishment during pine invasion of other vegetation types, the community and ecosystem shifts we document in our field data as a result of *B. pilularis* to *P. muricata* transition would have been greatly reduced without the presence of these fungi. Thus, while a recent review noted that no keystone mycorrhizal species had yet been identified (Banerjee et al. 2018), our evidence suggests that these ruderal, ectomycorrhizal fungi appear to function as keystone species, or perhaps a keystone guild, influencing the trophic structure of this system. While we lack the long-term data to illustrate these effects for symbiotrophic microbes associated with *B. pilularis* and *C. thyrsiflorus*, we expect the findings are generalizable to other systems. For example, while identifying symbiotrophic taxa is more difficult with bacteria where similar guild assignments are not possible, the long-term influence of N-fixing actinorhizal bacteria is evident in the soil environment even though they were not identified as abundant through our sequencing. There are relatively few examples of keystone microbial mutualists, but due to their relatively small biomass fractions, critical physiological impacts on plant growth (Friesen et al. 2011, Bennett et al. 2017), and ecosystem function (Vitousek and Walker 1989, Batterman et al. 2013, Clemmensen et al. 2013, Averill et al. 2014), we expect that many microbial mutualisms will fall into this category. Given their ability to find hosts during range expansion into non-host matrices (Hayward et al. 2015), this small set of ruderal symbionts may be particularly important agents of ecosystem change.

**Figure 6:**
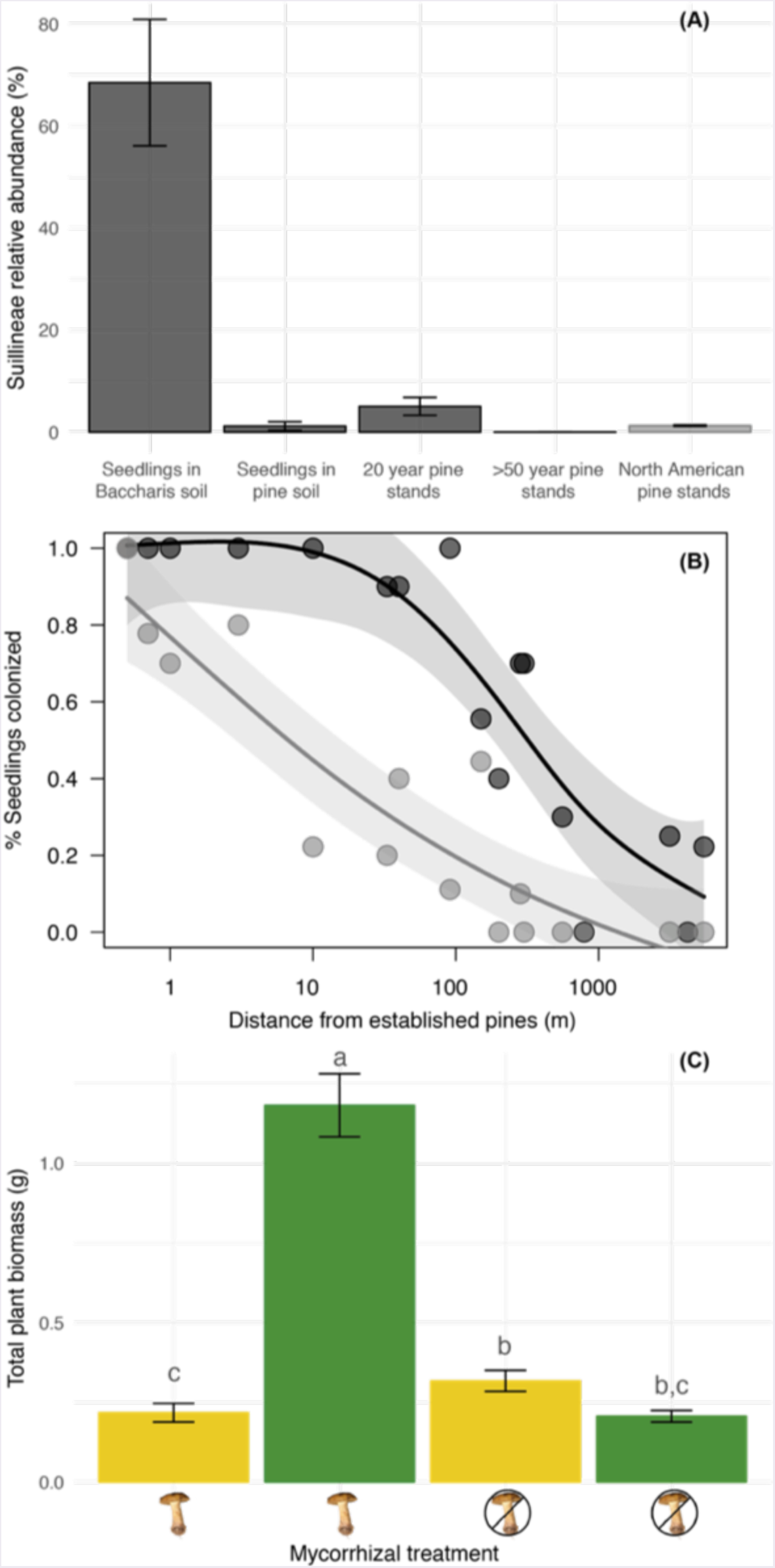
Fungi in the Suillineae are keystone mutualists. (a) Abundance of ectomycorrhizal fungi within the Suillineae (the genera *Rhizopogon* + *Suillus* at PRNS) in different habitats. Abundance is expressed as the proportion of ectomycorrhizal sequences detected in soils from bioassay seedlings and field plots in this study, a mature (>50 years) pine stand at PRNS, and from a continental study of pine soil communities across North America (from Talbot et al. 2014, Steidinger et al. in revision). (b) Suillineae are the primary ectomycorrhizal colonists away from established pines. The black lines shows the % of all pine seedlings colonized in a spatial bioassay of *Baccharis* soils collected at increasing distances from established pines. The grey line shows the fraction of seedlings that would be colonized in the absence of the Suillineae. The difference between the two lines shows that pine seedling colonization would be dramatically lowered in the absence of Suillineaea (data from Smith et al. 2018). (c) The absence of Suillineae puts pine seedlings at a competitive disadvantage. Figure shows biomass of *P. muricata* (green) and *B. pilularis* (yellow) seedlings in the presence and absence of spores from the ectomycorrhizal fungus *Suilluspungens* (data from Peay 2018).

## Conclusion

The evidence we have presented indicates that across a single landscape there may exist multiple, stable, ecosystem states within the soils. In our system, these states were determined by the dominant vegetation type at any given site, such that we observed a *Pinus*-associated ecosystem state, a *Baccharis*-associated state, and a *Ceonothus*-associated state in the PRNS soils. These alternative states are generated by feedbacks between plants and associated microbes, which together modify the local soil environment and the diverse community of organisms that inhabit it. While there has been much debate in the ecological literature about what constitutes alternative stable states (Connell and Sousa 1983), this system fulfills the primary conditions. First, these states developed from identical starting conditions (all were originally *B. pilularis*) and climate, thus the alternative trajectories we observe are not environmentally pre-determined. Second, these different states are persistent. Given that the patches of vegetation we sampled were >20 years old, these conditions have persisted for multiple generation from the perspective of soil microbes, which likely have generation times of weeks to years. From a plant perspective, while the landscape may be a shifting mosaic, all evidence suggests that these three species have coexisted for many generations (Forrestel et al. 2011). In early post-fire surveys at PRNS, Harvey and Holzman (2014) noted that plant communities were mixed initially but tended to become less diverse over time. The tendency to develop monodominant stands over time may result from positive feedbacks related to microbial communities or local environmental modification, as suggested by some theoretical models (Bever et al. 2010, Revillini et al. 2016).

Our study provides a holistic perspective on how ecosystems assemble. While the initial dispersal of plant species has a Gleasonian element of chance, their successful establishment requires the presence or timely co-dispersal of microbial symbionts. Symbiotic interactions take these microbial taxa from undetectable to dominant components of the microbial community, and in doing so increase likely increase the probability of host plant survival. Over time, the initial symbionts that facilitate host plant establishment are replaced by a greater influx of host associated symbionts and saprotrophic taxa responding to the new environmental conditions, ultimately creating a different soil ecosystem, but one that assembles predictably in much the way Clements imagined. While climate and soil fertility undoubtedly influence the structure and function of communities, our results show that within a given climate, biotic feedbacks with soil microbes can be sufficiently strong to produce discrete states with radically different properties. These effects are likely to be strongest in systems where dominant species host evolutionarily distinct forms of microbial symbionts (Vitousek and Walker 1989, Corrales et al. 2016), but subtle variation in the composition of saprotrophic microbes or within symbiotic guilds may also prove to be important.

## Acknowledgements

We thank Ben Becker and Point Reyes National Seashore for permission to conduct research. KGP received support from K.G.P. was supported by a Terman Fellowship from Stanford University, NSF Award DBI-1046115, and DOE Award DE-SC0016097

